# Immune infiltration is linked to increased transcriptional activity and distinct metabolic profiles in chordomas

**DOI:** 10.1101/2025.11.04.686078

**Authors:** Siddh van Oost, Rick Ursem, Zeynep B. Erdem, Sara Cardoso, Sanne Venneker, Ruud van der Breggen, Inge H. Briaire-de Bruijn, Alwine B. Kruisselbrink, Wilco C. Peul, Karoly Szuhai, Robert J.P. van der Wal, Rene van Zeijl, Bram Heijs, Noel F.C.C. de Miranda, Judith V.M.G. Bovee

## Abstract

Chordomas are ultra-rare cancers from the axial skeleton with limited treatment options. Chordomas can exhibit immunogenic features, and immune checkpoint blockade has demonstrated clinical activity in some patients, yet the determinants of these responses remain poorly understood. We aimed to elucidate the biological basis of immunogenicity in chordomas through spatial transcriptomic and metabolomic analyses.

We applied GeoMx digital spatial profiling to 15 chordomas, analyzing tumor and stromal compartments separately and comparing inflamed and non-inflamed tumors. Consecutive tissue sections were examined using MALDI-mass spectrometry imaging (MSI) at 20 µm spatial resolution. Findings were further explored through immunohistochemistry analyses. To assess whether transcriptional activity in tumor cells was associated with interferon-γ (IFN-γ) signaling, we tested the effect of IFN-γ treatment on chordoma cell lines.

Digital spatial profiling of inflamed and non-inflamed chordomas revealed striking differences in both the cancer cell and stromal compartments. Increased immune infiltration was strongly associated with elevated transcriptional activity in cancer cells. MALDI-MSI revealed distinct metabolic signatures and pronounced glycogen accumulation in inflamed chordomas. Nevertheless, IFN-γ stimulation of chordoma cell lines did not induce expression of *TBXT* (brachyury), the master transcriptional regulator of chordoma biology.

Collectively, our findings reveal a strong association between the transcriptomic and metabolic activity in chordomas and their immunogenicity. However, the causal direction of this relationship remains to be determined. Nevertheless, these findings are highly relevant for patient stratification, particularly in the context of immunotherapeutic strategies.

## Introduction

Chordomas are ultra-rare, malignant bone tumors demonstrating notochordal differentiation, arising almost exclusively in the axial skeleton, including the skull base, mobile spine and sacrum (1). The mainstay of treatment involves surgical resection and/or (proton beam) radiation (2, 3). While (proton beam) radiation can improve local control, *en bloc* resection remains the only curative option for localized disease. Clinical management is complicated by the proximity of tumors to critical neurovascular structures, often resulting in incomplete surgical margins, high rates of local recurrence, and high morbidity (2, 3). The lack of effective systemic therapies for advanced disease further emphasizes the urgent need for innovative and combinatory treatment strategies (4).

Intriguingly, T cell checkpoint blockade therapy has demonstrated promising results in a subset of patients, which suggests that chordomas harbor immunogenic features (5-7). However, biomarkers predicting responsiveness to T cell checkpoint blockade remain elusive. Previously, we shed light on natural immunity in chordomas through the identification of distinct immune contextures (8). Together with other studies (9-13), these findings suggest a dichotomy in chordoma inflammatory profiles, with some tumors showing pronounced immune infiltrate (inflamed), while others show little or no immune activity (non-inflamed). The mechanisms underlying this variability in immunogenic features remain unclear.

To investigate correlates of immunogenicity in chordomas, we applied an integrated spatial multimodal approach, combining spatial transcriptomics with spatial metabolomics. Inflamed tumors showed heightened transcriptional activity in both tumor and stromal compartments, suggesting a reciprocal biological activity between cancer cells and the immune microenvironment. Furthermore, inflamed chordomas displayed distinctive metabolic features. Collectively, these data highlight cancer cell activity as a potential driver of immunogenicity in chordomas, and could potentially inform patient stratification, particularly in the context of immunotherapy.

## Methods

### Patient samples

Snap-frozen tumor tissue was available for 18 samples from 15 patients with conventional chordoma (2 clival, 2 spinal and 11 sacrococcygeal), all of whom were part of a previously described cohort (8). An overview of the samples is provided in **Supplementary Table 1**. All samples included in this study were derived from surgical resection specimens, including three from patients who had received neoadjuvant radiotherapy.

### GeoMx digital spatial profiling

Slides were prepared in a cryostat and kept on dry-ice. Tissue sections of 5 µm were thaw-mounted onto pre-cooled slides to prevent RNA degradation and then stored at -80°C until use. Slide preparation was performed by the Utrecht Sequencing Facility (Utrecht, The Netherlands) according to NanoString’s protocol. In brief, slides were fixed using 10% neutral-buffered formalin, followed by overnight *in situ* hybridization using the Human NGS Whole Transcriptome Atlas (WTA) RNA (v1.0) probe set (NanoString Technologies). Slides were concurrently incubated with immunofluorescent antibodies, including pan-cytokeratin (AE-1/AE3, Novus Biologicals) to identify tumor cells and CD45 (2B11 + PD7/26, Novus Biologicals) to detect immune cells. Next, slides were stained with the GeoMx Nuclear Stain (Syto13, Thermo Fisher Scientific), after which slides were loaded into the GeoMx Digital Spatial Profiler instrument (NanoString Technologies) and scanned. For each sample, two to five rectangular regions of interest (ROIs) of variable sizes (up to 660×785um) were selected at the tumor-stroma border, representative of the overall tumor microenvironment. A total of 63 ROIs were UV-illuminated twice to capture the pan-cytokeratin segment separately from all other cells, which were subsequently classified as tumor and stroma respectively. CD45 staining was used to visualize immune cells and guide ROI selection, but not for segment capture.

### GeoMx data processing

All data processing and downstream analyses were performed in R (v.4.4.0) using the GeomxTools package (v.3.8.0). Quality control of the segments and the WTA probes was performed following NanoString’s analysis guideline. The used parameters and code are available on our lab’s Gitlab at https://git.lumc.nl/bstp/papers/spatial-profiling-of-chordomas. In short, no segments were excluded after quality control, but 21 probes were excluded after performing the Grubb’s test. A limit of quantification was calculated for the detected genes in each segment, after which genes were excluded that were detected in less than 5% (*n*=6) of all segments. This resulted in a final set of 8401 genes in 126 segments, with a minimum of 100 captured nuclei. The data was further processed with a Q3 quantile normalization and log2 transformation.

### GeoMx data analysis

Tumor segments were compared with stroma segments through a differential gene expression analysis. A linear mixed model (LMM) was used to correct for varying number of segments per sample, as described by NanoString’s analysis guideline. The results were visualized in a volcano plot with ggplot2 (R, v.3.5.1) and the top 20 differentially expressed genes (DEGs) were selected for annotation. All DEGs were visualized in a heatmap using ComplexHeatmap (R, v.2.20.0). Single-sample gene set enrichment analysis was performed using GSVA (R, v.1.52.3), which utilized an in-house curated WikiPathways gene list. The top differentially enriched biological pathways were identified comparing tumor and stroma segments through a similar LMM.

Next, segments were analyzed within their respective groups. Both tumor and stroma segments were clustered into two groups through graph-based clustering (Louvain algorithm) using bluster (R, v.1.14.0), which were then visualized using Uniform Manifold Approximation and Projection (UMAP). Differential gene expression analysis was again performed using an LMM, now comparing the identified clusters within their respective groups. Again, the top 20 DEGs per cluster were selected for visualization in a volcano plot. All DEGs were visualized per compartment in heatmaps. The number of captured nuclei and detected genes per segment were compared between the identified clusters using unpaired Student’s t-tests.

### T cell infiltration from imaging mass cytometry data

We previously analyzed samples from the current cohort by means of imaging mass cytometry (8). T cell counts from that analysis were extracted to validate the GeoMx classification of the samples into inflamed and non-inflamed categories. One sample (L5436) was excluded due to the imaging mass cytometry data including both conventional and dedifferentiated tumor components, while the frozen sample analyzed by GeoMx comprised only of conventional chordoma.

### MALDI-TOF-MSI

Concurrently with the preparation of the GeoMx slides, consecutive 5 µm tissue sections were cut from the frozen tissue samples, thaw-mounted onto conductive indium-tin-oxide (ITO)-coated glass slides (VisionTek Systems) and stored at -80°C until use. Before use, slides were placed in a vacuum freeze-drier for a minimum of 15 minutes.

Spatial metabolomics data was obtained using matrix-assisted laser desorption/ionization (MALDI)-MSI, using *N*-(1-naphthyl) ethylenediamine dihydrochloride (NEDC) (Sigma-Aldrich) as the matrix. A solution of 7 mg/mL NEDC in 70% MeOH (Actu-All Chemicals), 20% acetonitrile (Actu-All Chemicals) and 5% deionized water (MQ) was prepared. The tissue sections were coated with the matrix solution using a HTX M3+ sprayer (HTX Technologies). Detailed information about the matrix deposition step is available in **Supplementary Table 2**.

Spatial metabolomics data was acquired at a spatial resolution of 20 µm using a rapifleX MALDI-TOF/TOF system (Bruker Daltonics) operating in reflectron mode, negative polarity, and set to detect a mass range of 60 to 1200 *m/z*. Each pixel was acquired with 50 shots at a laser repetition rate of 10 kHz.

### High-resolution accurate-mass profiling acquisition

After spatial metabolomics data acquisition using the rapifleX MALDI-TOF/TOF, each tissue section was also subjected to high-resolution, accurate-mass (HRAM) profiling using a 15T solariX MALDI-FTICR mass spectrometer (Bruker Daltonics). The MALDI-FTICR was operated in negative-ion mode, using a 2M transient length, and set to detect a mass range of 400 to 1000 *m/z*. Each HRAM lipid/metabolite profile was collected from 25 laser shots, using a laser repetition rate of 120 Hz and with the laser focus set to ‘minimum’. After visual inspection of the spatial metabolomics results, HRAM profiles were acquired from regions that exhibited distinct lipid and metabolite signatures. After completing the spatial metabolomics and HRAM profiling, residual MALDI matrix was removed from the slides by ethanol washing and rehydration, followed by hematoxylin and eosin (H&E) staining according to standard diagnostic procedures. Finally, high resolution optical images of the H&E-stained tissue sections were recorded using a Zeiss scanner.

### MALDI-TOF-MSI data processing

All spatial metabolomics data were imported into SCiLS Lab (v.2023b 11.01.14623, Bruker Daltonics) and subjected to baseline correction using the TopHat algorithm (width 200). The average spectrum of all tissues was exported from SCiLS Lab and used for peak picking in mMass (v.5.5.0) with a Signal-to-Noise Ratio (SNR) > 9, followed by removal of matrix derived peaks by image correlation to the principal matrix peak ([NEDC+Cl]^-^). Bisecting *k*-means clustering was employed in SCiLS Lab to differentiate on-tissue pixels from off-tissue pixels and areas without signal. Total Ion Count (TIC) normalized peak intensities and masses were exported as .csv files and processed further. Masses were internally calibrated to a list of abundant lipids and molecules commonly detected as negative ions when using NEDC as the matrix (**Supplementary Table 3**). Spectra were normalized to the total ion count before further analysis. For small metabolite assignment, MALDI-TOF-MSI peaks were matched to the HMDB database (accessed 10/07/2024) with a mass error tolerance of ± 30 ppm allowing for [M-H]^-^ and [M+Cl]^-^ adducts.

### Lipid/metabolite assignment based on high-resolution accurate-mass profiling

The HRAM profiles obtained from the MALDI-FTICR mass spectrometer were imported and processed in R (v.4.1) with MALDIquantForeign (v.0.13) and MALDIquant (v.1.20). Spectra were aligned, peak picked, and internally calibrated similarly to the MSI data (**Supplementary Table 4**).

Resulting peaks were matched to the LIPIDMAPS LMSD database (accessed 14/01/2021) for phosphatidic acid (PA), phosphatidylethanolamine (PE), phosphatidylinositol (PI), phosphatidylserine (PS), and phosphatidylglycerol (PG), using a mass tolerance of ± 3 ppm and allowing for [M-H]^-^ and [M+Cl]^-^ adducts. This list was transferred to the lower mass resolution spatial metabolomics MALDI-TOF-MSI data, matching every MALDI-TOF-MSI peak with its analogous peak in the HRAM MALDI-FTICR spectra, using a tolerance of ± 40 ppm. Finally, all lipid assignments were manually curated to remove false positives and to ensure that all mass spectral peaks, which were used for the subsequent data analysis steps, corresponded to a single lipid species in the HRAM lipid profiles.

### MALDI-TOF-MSI data analysis

Downstream analysis of the curated spatial metabolomics data was performed using Seurat (R, v.4.2). Per sample, data points (pixels) corresponding to holes in the tissue or necrotic tissue were removed after *k*-means clustering and subsequent clustering with Seurat. The lipid/metabolite abundance data of all samples was transformed into a single Seurat object followed by log normalization, after which the data was clustered (Louvain algorithm) and visualized by a dimensionality reduction using UMAP. For differential analyses, clusters were classified as tumor or stroma by comparing their spatial localization with corresponding histopathological images, after which the mean lipid/metabolite intensity per region was calculated per sample.

### Immunohistochemistry

Immunohistochemical detection for Ki-67 (RTU, MIB-1, 1:1, Dako, GA626) was performed as described previously (8). Formalin-fixed paraffin-embedded (FFPE) tissue blocks corresponding to the same tumors used for the GeoMx/MALDI analyses were assessed. Given its overall low expression, Ki-67^+^ histological tumor regions were selected in QuPath (v.0.3.1) to quantify the percentage of Ki-67^+^ tumor cells per 1000 tumor cells.

### Interferon-γ treatment of chordoma cell lines

One chordoma cell line from a primary tumor was established in-house (L2895) and additional chordoma cell lines were acquired via the Chordoma Foundation, including MUG-CHOR1, UCH2 and UM-CHOR1. All chordoma cell lines were cultured in IMDM/RPMI 1640 (4:1, Gibco) supplemented with 10% non-inactivated FBS (F7524, Sigma) and 1% Insulin-Transferrin-Selenium (ITS-G, Gibco). Cells were seeded in 50 µg/mL Collagen Type I (50202, Ibidi) coated 6-well plates at a cell density of 50.000 cells/well. After 11 days, cells were either not treated (mock) or treated with 100 U/mL IFN-γ (11343536, Immunotools). After one or three days of treatment, cells were harvested and pellets were stored at -80°C until RNA isolation was performed.

### qPCR

Total RNA was isolated from the cell pellets using the NucleoSpin RNA Mini kit (Macherey-Nagel), and cDNA synthesis was performed using the Transcriptor First Strand cDNA Synthesis Kit (Roche). Quantitative PCR (qPCR) was performed to quantify four genes: *TBP* and *HPRT1* as housekeeping genes (14, 15), *B2M* as positive control and *TBXT* as gene of interest. Transcript structure data from Ensembl indicates that *TBXT* (ENSG00000164458) encodes three protein-coding isoforms: *TBXT-201* (ENST00000296946) and *TBXT-203* (ENST00000366876) both include exon 6, while *TBXT-202* (ENST00000366871) excludes exon 6 (16). Isoform-specific primers were therefore designed to distinguish between the “long” isoforms (*TBXT-201/203*) and the “short” isoform (*TBXT-202*), as well as primers targeting a region common to all *TBXT* transcripts (*TBXT* (all transcripts)). Primer sequences are provided in **Table 1**. The qPCR reactions were performed using iQ SYBR Green Supermix (Bio-Rad) and run on a C1000 Touch Thermal Cycler equipped with the CFX384 Touch Real-Time PCR Detection System (Bio-Rad). A standard PCR protocol was followed by annealing/extension at 59.2 °C for 45 seconds. All reactions were carried out in triplicate. Reactions were performed in 384-well Hard-Shell thin-wall PCR plates (Bio-Rad, HSP3805) sealed with Microseal ‘B’ adhesive film (Bio-Rad, MSB1001). Data were analyzed using CFX Maestro software (v.5.3.022.1030, Bio-Rad).

**Table 1.**
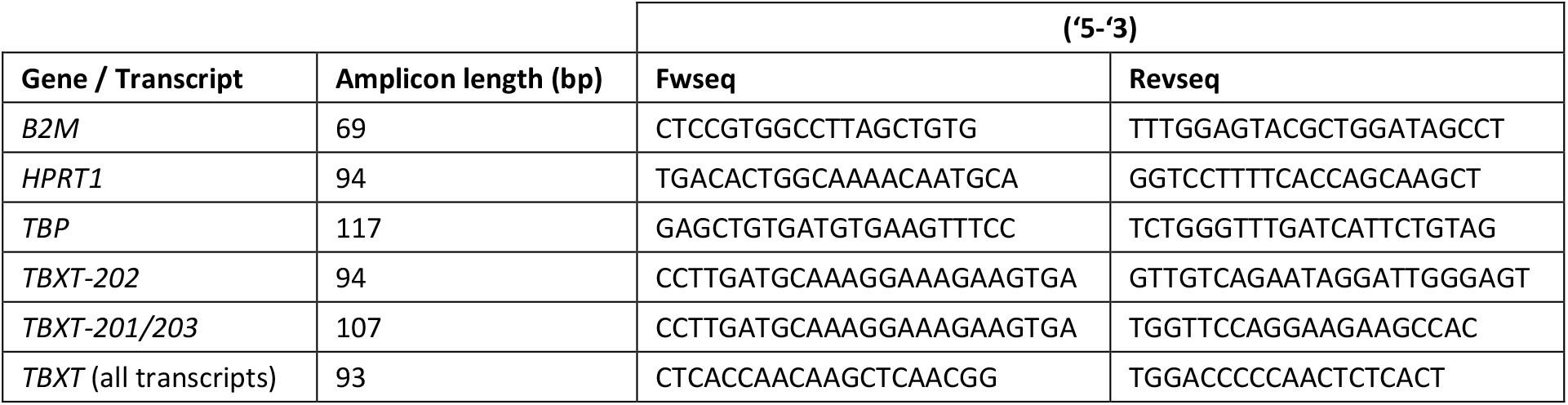
Primer sequences. Fwseq = forward primer sequence; Revseq = reverse primer sequence; bp = base pairs.

### qPCR data analysis

Average Ct values were calculated from technical triplicates. For each condition, ΔCt values were determined by subtracting the Ct of the housekeeping gene from that of the gene or transcript of interest. ΔΔCt values were then calculated by subtracting the ΔCt of the control condition (mock) from the ΔCt of the condition of interest. Fold changes in expression were computed using the 2^−ΔΔCt^ method. These calculations were performed separately using *HPRT1* and *TBP*, after which the average and standard deviation were calculated across both housekeeping genes. Boxplots were generated using ggplot2 (R, v.3.5.1) to visualize the normalized fold change in expression of the target genes and transcripts.

### PAS staining

FFPE tissue sections from corresponding tumors were stained for Periodic acid–Schiff (PAS) and diastase-PAS (dPAS) staining. PAS staining was performed using the Artisan PAS Stain Kit (AR165, Artisan, Agilent), while dPAS staining was carried out using the same kit in combination with alpha-amylase digestion using the Artisan Alpha-Amylase reagent (AR171, Artisan, Agilent). All staining procedures were executed on the Artisan Link Pro Special Staining System (Dako, Agilent) according to the manufacturer’s protocols. Comparison of PAS and dPAS staining demonstrated that the majority of the PAS signal in tumor cells was attributable to glycogen, as diastase digestion abolished most of the staining (**Supplementary Figure 1**). Samples were subsequently classified as high or low based on the relative PAS staining intensity within the tumor cell compartment, as assessed by one observer (ZBE). The resulting groups were visualized in cumulative bar plots per identified GeoMx cluster using ggplot2 (R, v.3.5.1).

## Ethical approval

Samples were pseudo anonymized and handled according to the ethical guidelines outlined in ‘Code for Proper Secondary Use of Human Tissue in The Netherlands’ by the Dutch Federation of Medical Scientific Societies. A waiver of consent was obtained from the medical ethical evaluation committee (Medisch-Ethische Toetsingscommissie Leiden Den Haag Delft; protocol number: B17.036 and B20.067). Therefore, this study adheres to the Declaration of Helsinki.

## Data availability

The utilized imaging mass cytometry data is available as raw data in BioStudies at S-BIAD830 or as processed data on the LUMC Bone & Soft Tissue Pathology Group GitLab (https://git.lumc.nl/bstp/papers/multimodal-profiling-of-chordomas). The raw and processed GeoMx digital spatial profiling data and the processed MALDI-MSI data generated in this study are publicly available on Figshare (https://figshare.com/s/4d52d0129b9e0f9c7a8c). The R code used to process the data from this study is also published on our group’s GitLab (https://git.lumc.nl/bstp/papers/spatial-profiling-of-chordomas). All other data are available from the corresponding author upon reasonable request.

## Author contributions

SvO contributed to the conceptualization, funding acquisition, data curation, software, formal analysis, validation, investigation, visualization, methodology, writing-original draft, project administration, and writing-review editing. RU contributed to the data curation, software, formal analysis, validation, investigation, visualization, methodology, writing-original draft, and writing-review editing. ZBE contributed to the validation, investigation, and writing-review editing. SC contributed to the validation, methodology and writing-review editing. SV contributed to the validation, investigation, methodology, writing-original draft, and writing-review editing. RvdB contributed to the formal analysis, validation, investigation, methodology, and writing-review editing. IHBdB contributed to the investigation, methodology and writing-review editing. ABK contributed to the validation, investigation, methodology, writing-original draft, and writing-review editing. WCP contributed to the resources and writing-review editing. KS contributed to resources, writing-review editing. RJPvdW contributed to the resources and writing-review editing. RvZ contributed to the investigation and writing-review editing. BH contributed to the software, formal analysis, investigation, methodology, and writing-review editing. NFCCdM contributed to the conceptualization, supervision, funding acquisition, investigation, methodology, writing-original draft, and writing-review editing. JVMGB contributed to the conceptualization, supervision, funding acquisition, investigation, methodology, writing-original draft, and writing-review editing.

## Declaration of generative AI and AI-assisted technologies in the writing process

During the preparation of this work the author(s) used ChatGPT in order to improve the readability and language of the manuscript. After using this tool/service, the author(s) reviewed and edited the content as needed and take(s) full responsibility for the content of the published article

## Acknowledgements

We acknowledge the Utrecht Sequencing Facility (USEQ) for providing sequencing service and data. USEQ is subsidized by the University Medical Center Utrecht and The Netherlands X-omics Initiative (NWO project 184.034.019).

## Results

### Inflammation is associated with increased transcriptional activity

To investigate transcriptional profiles of both cancer and stromal compartments that associate with immunogenic features in chordomas, GeoMx was performed on frozen sections from 15 conventional chordomas. For each sample, two to five ROIs along the tumor-stroma border were selected (**Supplementary Figure 2**). Gene expression profiles were then delineated into tumor and stroma compartments based on pan-cytokeratin expression. Differential gene expression analysis was performed to compare tumor segments with stroma segments (**Figure 1a, Supplementary Figure 3**). This analysis confirmed the enrichment of chordoma-associated genes in tumor segments, including

**Figure 1.**
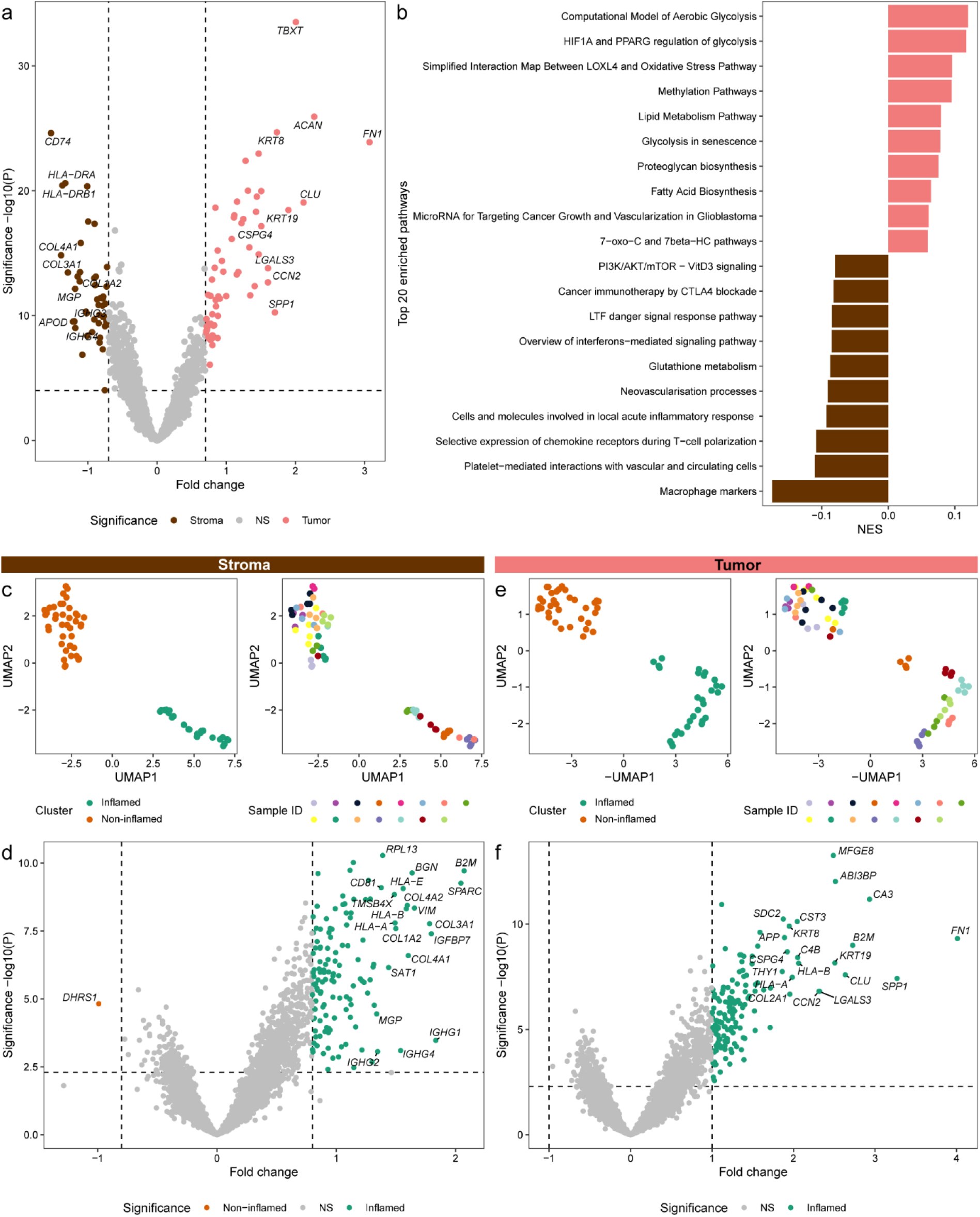
Inflammation is associated with increased transcriptional activity. a) Volcano plot presenting the differential gene expression between stroma (brown) and tumor (pink) segments. The top 10 differentially expressed genes (DEGs) per segment, ranked by fold change, are highlighted. Statistical significance was determined using a linear mixed model (LMM) correcting for sample ID. b) Bar plot of the normalized enrichment score (NES) for the top 10 differentially enriched pathways per segment type, based on a curated WikiPathways list. Statistical significance was assessed using an LMM correcting for sample ID. c) UMAP visualizations of stroma segments using the top 20% most variably expressed genes. Clusters were defined via graph-based clustering. Segments are colored by cluster (left) and sample ID (right). d) Volcano plots comparing non-inflamed (orange) versus inflamed (green) stroma segments. The top 20 DEGs are highlighted per cluster. Statistical significance was determined using an LMM correcting for sample ID. e) UMAP visualizations of tumor segments using the top 20% most variably expressed genes. Clusters were defined via graph-based clustering. Segments are colored by cluster (left) and sample ID (right). f) Volcano plots comparing non-inflamed (orange) versus inflamed (green) tumor segments. The top 20 DEGs are highlighted per cluster. Statistical significance was determined using an LMM correcting for sample ID.

*TBXT, ACAN, KRT8* and *KRT19*, and immune- and stroma-related genes in stroma segments, such as genes encoding collagens, Human Leukocyte Antigen class II molecules, or immunoglobulins (**Figure 1a, Supplementary Figure 3**). As expected, many genes which have been described as targets of *TBXT* (e.g. *CD109, MRGPRX3, SOX9*) (17) were also enriched in the tumor segments. Next, biological pathways were explored through single-sample gene set enrichment analysis for all segments using a manually curated WikiPathways list. Accordingly, differential pathway analysis comparing tumor with stroma segments revealed that tumor segments were enriched for cancer-associated pathways, such as glycolysis, while stroma segments showed enrichment of immune- and stroma-related pathways like inflammatory signaling and neovascularization (**Figure 1b**).

Following this general characterization, tumor and stroma segments were interrogated separately to investigate the association between cancer cell-specific gene expression and immune infiltration. Using the top 20% most variably expressed genes (1680 genes), graph-based clustering identified two distinct groups of stroma segments, which were visualized with UMAP (**Figure 1c**). Differential gene expression analysis indicated that these clusters reflected inflamed and non-inflamed stroma segments due to the enrichment of immune-related genes or the lack thereof (**Figure 1d, Supplementary Figure 4**). These genes indicated enrichment of (plasma) B cells (*IGHG1, IGHG4*), antigen presenting cells (*HLA-DRA, HLA-DQB1*), macrophages (*CD68, LYZ*), fibroblasts (*COL1A2, COL3A1*), endothelial cells (*ACTA2, COL4A1*), complement (*C1R, C4B*), and T cells (*TRAC*). This pattern was accompanied by increased numbers of captured nuclei and detected genes in inflamed segments (**Supplementary Figure 5a+b**), reflecting increased cell densities of immune and stromal cells in these segments.

Building on this clear dichotomy within the stroma segments, tumor segments were also clustered into two groups using graph-based clustering of the top 20% most variably expressed genes and visualized with UMAP (**Figure 1e**). Interestingly, these tumor clusters closely mirrored the inflammation-associated pattern observed in the stroma clusters, with 14 of the 15 samples belonging to the same group across both compartments. Supporting these observations, previously generated imaging mass cytometry data from the same tumors showed increased T cell infiltration in the inflamed tumor cluster compared with the non-inflamed cluster (**Supplementary Figure 5c**). As expected, genes associated with immunity (*B2M, C1S, C4B, HLA-A, HLA-B, HLA-E, IFITM3*) had higher expression in the inflamed tumor segments than in the non-inflamed segments (**Figure 1f, Supplementary Figure 6**). More intriguingly, however, several chordoma-associated genes (*TBXT, CA3, CCN2, CD24, COL2A1, KRT19, SOX9*) also had higher expression in the inflamed tumor segments than in the non-inflamed segments (**Figure 1f, Supplementary Figure 6**). Interestingly, unlike in the stromal compartment, the number of captured nuclei did not differ between inflamed and non-inflamed tumor segments (**Supplementary Figure 5d**), while the number of detected genes was higher in the inflamed segments compared to the non-inflamed segments (**Supplementary Figure 5e**). Together with the overall enrichment of DEGs in the inflamed segments, these results indicate that chordoma cells in inflamed regions exhibit higher transcriptional activity compared to those in non-inflamed regions.

### Inflammation is associated with distinct lipid and metabolic profiles

To explore the metabolic and lipid profiles of chordomas, we performed MALDI-MSI at 20 μm resolution on whole slides from consecutive frozen sections of the same samples analyzed by GeoMx. For the data-driven annotation of the tissues, all measured lipid profiles were visualized using UMAP (**Figure 2a)** and, following clustering, were mapped back to their original spatial coordinates. By aligning with histopathological images, the lipid clusters enabled differentiation between tumor and stromal regions (**Figure 2b)**. This facilitated comparative analyses both between tumor and stroma as well as between inflamed and non-inflamed regions within each compartment. Following classification of the lipid clusters as either tumor- (clusters 1-10) or stroma-associated (cluster 0), tumor regions exhibited strong patient-specific profiles, while stromal profiles were relatively similar across patients (**Figure 2a+b**). Differential analysis showed that tumor areas were enriched for a diverse set of lipids, predominantly phosphatidylinositol (PI) species, along with some lysophosphatidylethanolamines (LPE), phosphatidic acid (PA), phosphatidylethanolamine (PE), and phosphatidylserine (PS) species, while stromal regions were enriched for only a single PS species (**Figure 2c**). Notably, tumors were enriched in polyunsaturated lipids (with more than two double bonds), suggesting more flexible and dynamic cell membranes. In contrast, the PS species enriched in stroma (PS(36:1)) is more saturated, containing only a single double bond, consistent with more rigid and stable membranes.

**Figure 2.**
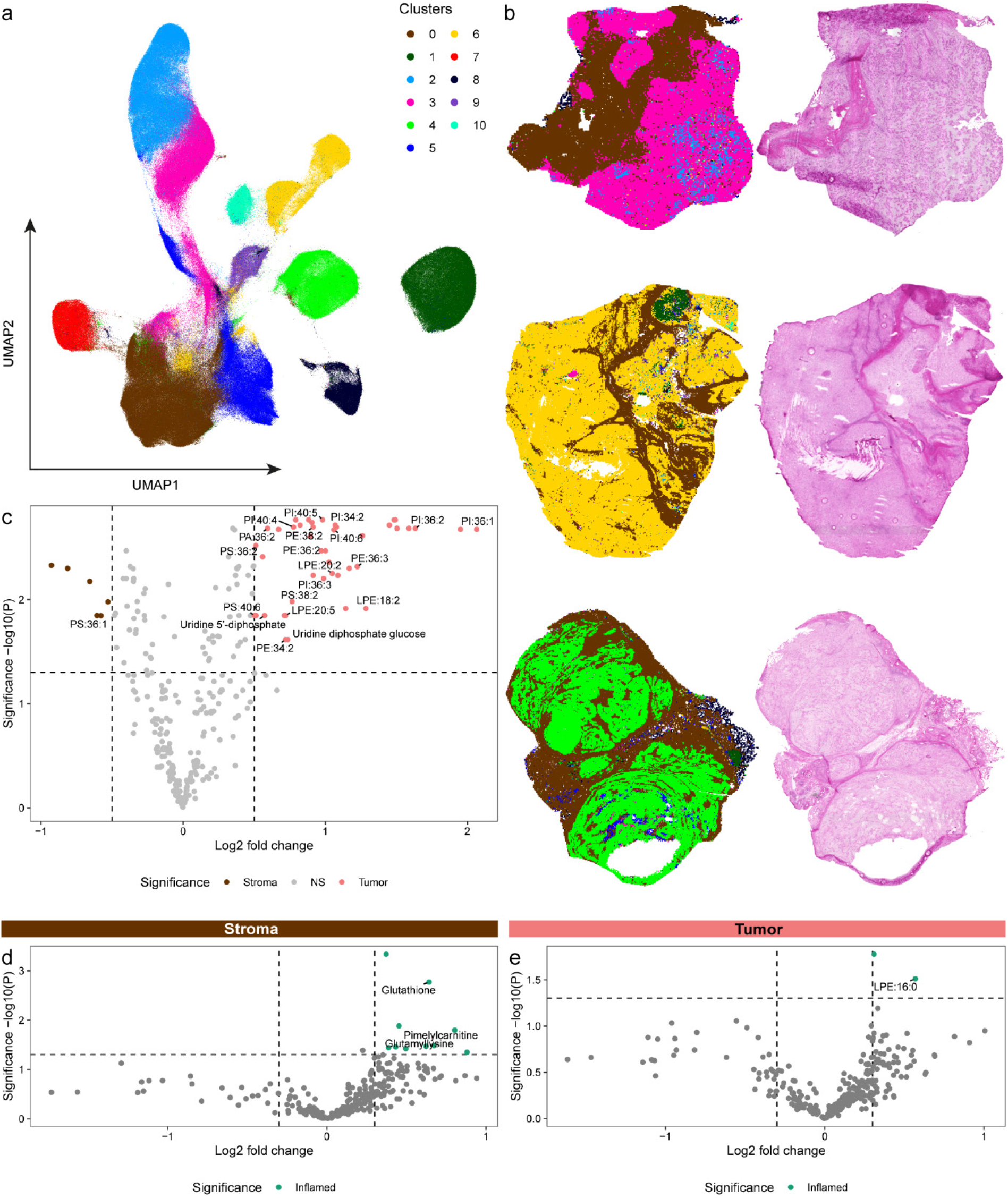
Spatial lipid and metabolite profiles of chordoma. a) UMAP visualization of all measured lipid profiles from the MALDI-MSI data. Clusters were identified through Louvain clustering. b) Spatial visualization of the identified clusters (stroma = brown, tumor = pink, yellow, green) next to corresponding histopathological images for 3 samples. c) Volcano plot presenting the differential lipid/metabolite abundance analysis comparing tumor with stroma regions. Significance was assessed by paired Student’s t-tests, which were corrected for multiple testing with the Benjamini-Hochberg method. d+e) Volcano plots presenting the differential lipid/metabolite abundance analysis comparing inflamed with non-inflamed samples, for tumor regions (d) and for stromal regions (e). Significance was determined by unpaired Student’s t-tests.

Next, we compared inflamed and non-inflamed tumors, as defined by the GeoMx analysis, using differential analysis within each compartment (**Figure 2d+e**). The resulting profiles mirrored the GeoMx findings, albeit less pronounced: inflamed chordomas displayed a greater number of differentially abundant lipids and metabolites in both tumor and stromal regions than non-inflamed chordomas. In particular, glutathione was enriched in the inflamed stroma, while the saturated lipid species LPE(16:0) was elevated in inflamed tumor regions. Together, these findings show that inflamed chordomas exhibit distinct lipid and metabolic profiles compared with non-inflamed tumors.

### Inflammation is associated with increased glycogen accumulation and proliferation

The MSI analysis comparing tumor with stroma revealed that the metabolite uridine diphosphate (UDP)-glucose, the key substrate for glycogen synthase, was enriched in tumor regions compared to stroma (**Figure 2c**). Consistently, the GeoMx analysis showed increased expression of two glycogen synthesis–related genes, *PPP1R3C* and *UGP2*, in inflamed tumor segments (**Supplementary Figure 6**). *PPP1R3C* encodes a subunit of the protein phosphatase 1 (PP1) complex, which activates glycogen synthase while inhibiting glycogen phosphorylase, thereby promoting glycogen storage. *UGP2* encodes UDP-Glucose Pyrophosphorylase 2, an enzyme that catalyzes the conversion of glucose-1-phosphate and uridine triphosphate (UTP) into UDP-glucose. Supporting these findings, PAS/dPAS staining demonstrated abundant glycogen accumulation in tumor cells of most chordomas, with inflamed tumors generally exhibiting stronger staining intensity and greater glycogen abundance than non-inflamed tumors (**Figure 3a-c**). Collectively, these results indicate that glycogen synthesis pathways are active in chordomas, and that inflamed tumors exhibit increased glycogen accumulation.

**Figure 3.**
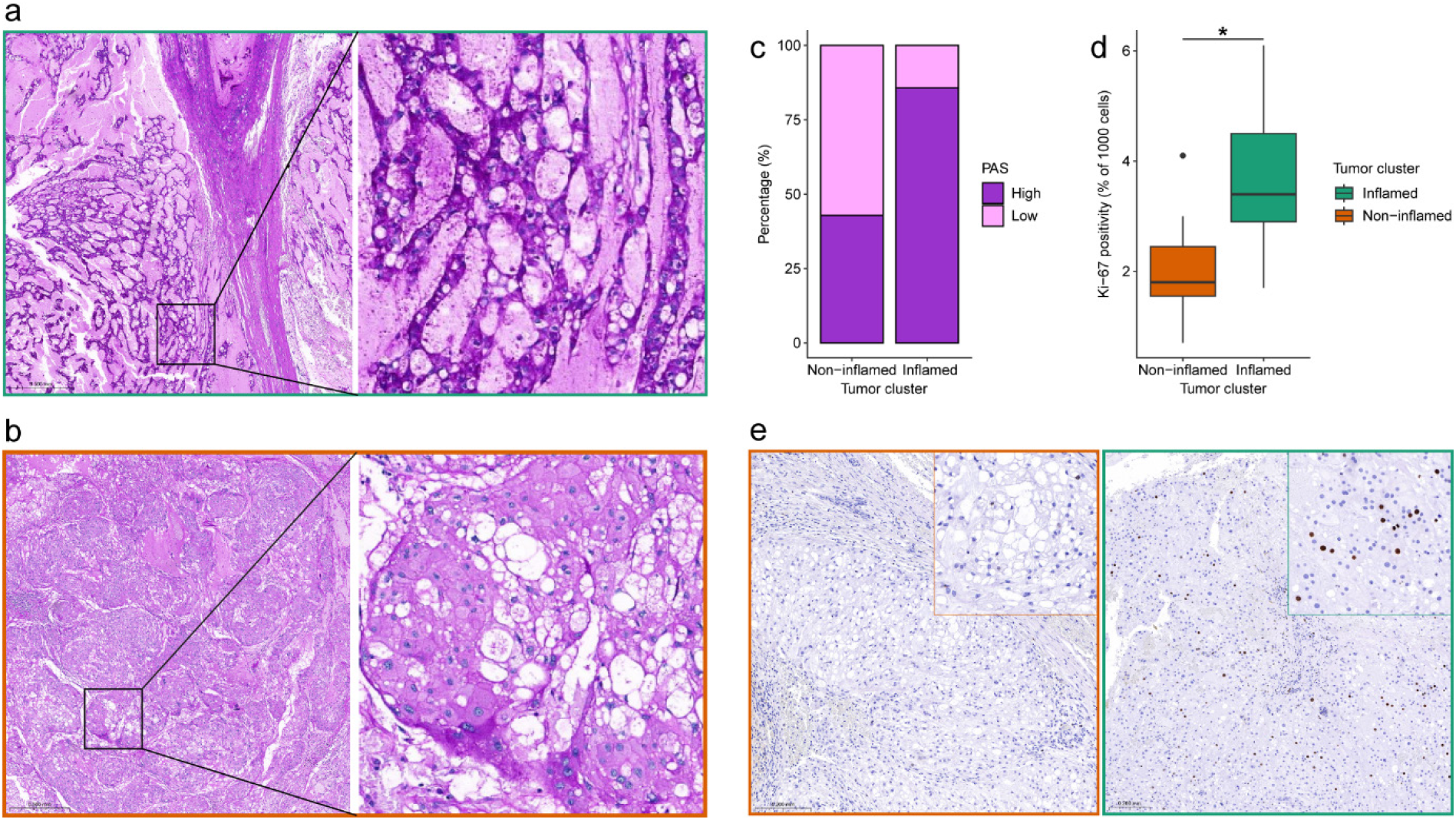
Glycogen accumulation and proliferation are increased in inflamed chordomas. a+b) Example images of the PAS staining of an inflamed chordoma (a), classified as high PAS staining, and a non-inflamed chordoma (b), classified as low PAS staining. c) Bar plots displaying the percentage of chordomas with high and low PAS staining, grouped per tumor cluster. d) Boxplots presenting the percentage of Ki-67^+^ tumor cells in proliferative hotspot areas, grouped per tumor cluster. Statistical significance was assessed through a Student’s t-test: * = *P* < 0.05. e) Example of Ki-67 immunohistochemistry showing Ki-67 expression in a non-inflamed (left) and inflamed (right) chordoma.

Following this, we examined whether increased transcriptional activity and glycogen accumulation were associated with increased proliferation of tumor cells. Although proliferation-associated genes were not detected in the GeoMx dataset, likely due to their low expression and the limitations of the WTA, immunohistochemistry for Ki-67 showed that inflamed tumors were more proliferative than the non-inflamed tumors (**Figure 3d+e**). Nonetheless, proliferation remained very low compared with that observed in carcinomas or high-grade sarcomas, with only 2 samples showing Ki-67 positivity above 5%.

### Interferon-γ does not induce *TBXT* expression in chordoma cell lines

To determine whether the heightened transcriptional activity of inflamed tumors was driven by IFN-γ — given that several IFN-γ targets (e.g., HLA class I genes) were overexpressed by tumor cells in these tumors — we tested whether IFN-γ could induce *TBXT* expression. Multiple chordoma cell lines were treated with IFN-γ, and expression of *B2M* (positive control) and *TBXT* were assessed by qPCR. Ct values were normalized to two established chordoma housekeeping genes, *HPRT1* and *TBP* (14, 15), both of which showed stable expression across treatment conditions (**Supplementary Figure 7**). As expected, *B2M* expression increased in all cell lines after treatment with IFN-γ (**Figure 4**). In contrast, the expression of *TBXT* isoforms remained unchanged, indicating that *TBXT* is not regulated by IFN-γ·

**Figure 4.**
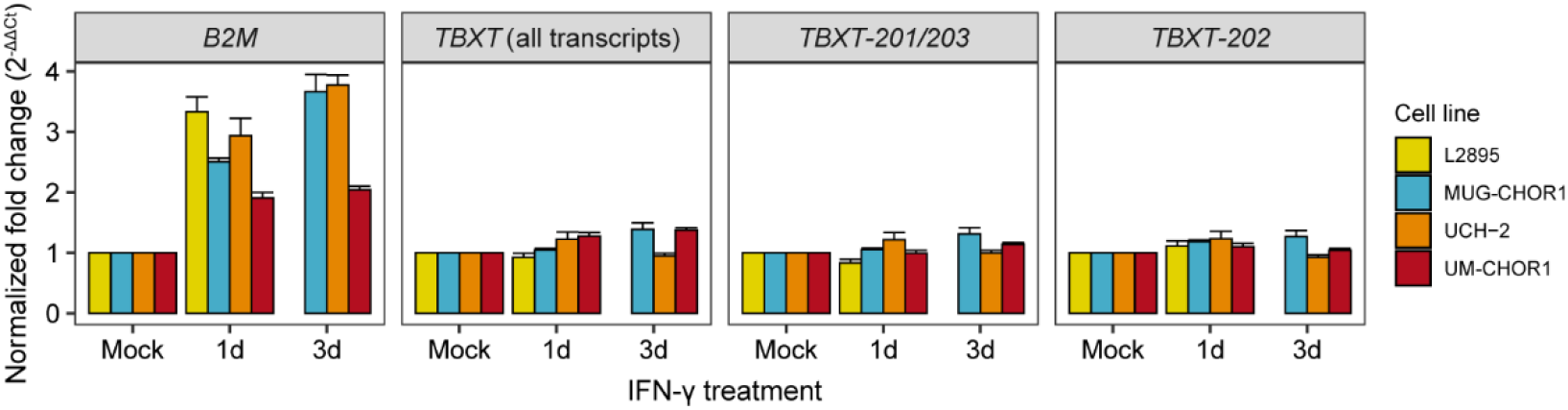
*TBXT* expression is not regulated by IFN-γ. Normalized expression fold change for each chordoma cell line and treatment condition, for *B2M* and the *TBXT* isoforms. All values were normalized for both housekeeping genes, *HPRT1* and *TBP*. Abbreviations: 1d = 1 day; 3d = 3days.

## Discussion

In this study, we applied an integrated spatial approach to explore correlates of immunogenicity in chordoma. Our findings revealed a clear dichotomy between inflamed and non-inflamed chordomas regarding overall transcriptional activity across both tumor and stromal compartments. Inflamed tumors exhibited higher expression of immune-related genes, such as those encoding HLA class I molecules and complement factors, as well as chordoma-associated genes, including *TBXT, SOX9*, and *COL2A1*, within the tumor cell compartment. This elevated transcriptional activity was accompanied by distinct metabolic profiles and increased glycogen accumulation, suggesting an overall heightened tumor cell activity state in inflamed tumors. To examine whether inflammatory signaling could contribute to this transcriptional activity, we investigated whether IFN-γ was able to induce the master regulator of chordoma biology, *TBXT*. While chordoma cells were responsive to IFN-γ, IFN-γ signaling did not impact *TBXT* levels, indicating that IFN-γ signaling alone is unlikely to explain the elevated *TBXT* levels observed in inflamed tumors. Nevertheless, other cytokines and mediators released by infiltrating immune or stromal cells may still influence tumor cell transcriptional activity. Future studies should investigate the roles of additional cytokines and tumor– immune–stroma interactions in shaping the chordoma transcriptome.

These findings extend prior work, including our own, that identified distinct immune contextures in chordoma (8-13), and highlight a strong association between tumor cell transcriptional activity and immune infiltration. Although the causal link remains unresolved, differences in *TBXT* expression, potentially shaped by underlying genetic or epigenetic variation, could help explain the heterogeneity observed in tumor phenotypes and immune infiltration. For instance, duplications or amplifications of the *TBXT* locus have been reported in 15-30% of chordoma patients (18-20). In addition, chordomas are transcriptionally addicted to super-enhancer–driven *TBXT* expression (21-23), and variation in enhancer strength or chromatin state between tumors could potentially contribute to the variable expression levels. Because *TBXT* regulates a broad range of transcriptional programs (17, 24), including cancer-associated pathways, its differential activity could indirectly influence pathways related to antigen processing or cellular stress responses, although this remains to be established experimentally.

In addition to its role as a master regulator, *TBXT* has been identified as an immunogenic antigen in chordoma. Brachyury-derived peptides have been shown to expand T cells *in vitro* (25, 26), and vaccination strategies targeting brachyury have elicited *TBXT*-specific T cell responses in a subset of chordoma patients (27, 28). Furthermore, a recent immunopeptidomics reported naturally presented HLA peptides derived from brachyury across common alleles, covering 80% of their patient population (29). Together, these findings underscore the immunogenic potential of *TBXT* and support further investigation into how its expression variability and peptide presentation contribute to immune recognition and response in chordoma.

Our spatial metabolomics analysis revealed that tumor regions of chordoma were enriched in polyunsaturated lipids. Because these lipids contain multiple double bonds, they are particularly susceptible to lipid peroxidation – a process in which reactive oxygen species attack fatty acids, generating toxic radicals that compromise membrane integrity and can trigger ferroptosis, a regulated form of cell death (30). Consistent with this, the Chordoma Foundation recently reported preclinical evidence indicating a ferroptosis vulnerability in chordoma cell lines (31), reinforcing our observations. In line with this concept, inflamed tumors were enriched for a saturated LPE species (16:0) in tumor regions and glutathione in stromal regions. Glutathione, a key antioxidant, and saturated lipids, which are less prone to peroxidation, suggest coordinated adaptations to oxidative stress. These findings suggest that inflamed tumors experience increased oxidative stress, potentially driven by heightened tumor cell activity, increased immune infiltration, or both (32). Further studies will be required to elucidate how oxidative stress interacts with tumor cell activity and immune infiltration in chordoma.

In addition to lipid and antioxidant adaptations, inflamed chordomas also exhibited increased glycogen accumulation, accompanied by higher expression of glycogen synthesis–related genes. Glycogen serves as a critical energy reserve that supports cell survival under hypoxic conditions (33). Its accumulation in inflamed tumors may therefore provide a metabolic advantage that sustains cellular activity and viability within an immune-infiltrated, hypoxic microenvironment. Glycogen storage is, however, a recognized feature of chordomas (34), likely associated with their notochordal differentiation (35). This notion is supported by the observation that *PPP1R3C* is a transcriptional target of *TBXT* (17). Accordingly, the increased glycogen accumulation observed in inflamed chordomas could also result from enhanced *TBXT* activity rather than representing a purely microenvironmental adaptation.

Going forward, it will be important to identify proxies or biomarkers that reflect tumor cell activity or immunogenic potential, enabling broader clinical translation. Notably, recent work by Crombé *et al*. demonstrated that high ^18F-FDG uptake correlates with increased immune cell densities in high-grade soft tissue sarcomas (36). Consistent with our findings, this suggests that metabolic imaging approaches such as FDG-PET could potentially serve as non-invasive tools to identify immunogenic chordoma subsets and guide patient stratification for immunotherapeutic strategies. This concept, however, will require validation in larger, ideally multi-institutional, cohorts to confirm its clinical utility.

In conclusion, our integrated spatial approach demonstrates that inflamed chordomas exhibit heightened transcriptional activity and distinct metabolic signatures, including increased glycogen accumulation. These features underscore a close interplay between tumor metabolism, transcriptional activity, and immune infiltration. While the causal relationships among these processes remain to be established, they point to metabolic an transcriptional states as potential determinants or indicators of tumor immunogenicity. Ultimately, such integrated molecular and spatial insights may guide the development of biomarkers and inform patient stratification for future immunotherapeutic strategies.

