## Supplementary Data for "Immune infiltration is linked to increased transcriptional activity and distinct metabolic profiles in chordomas"

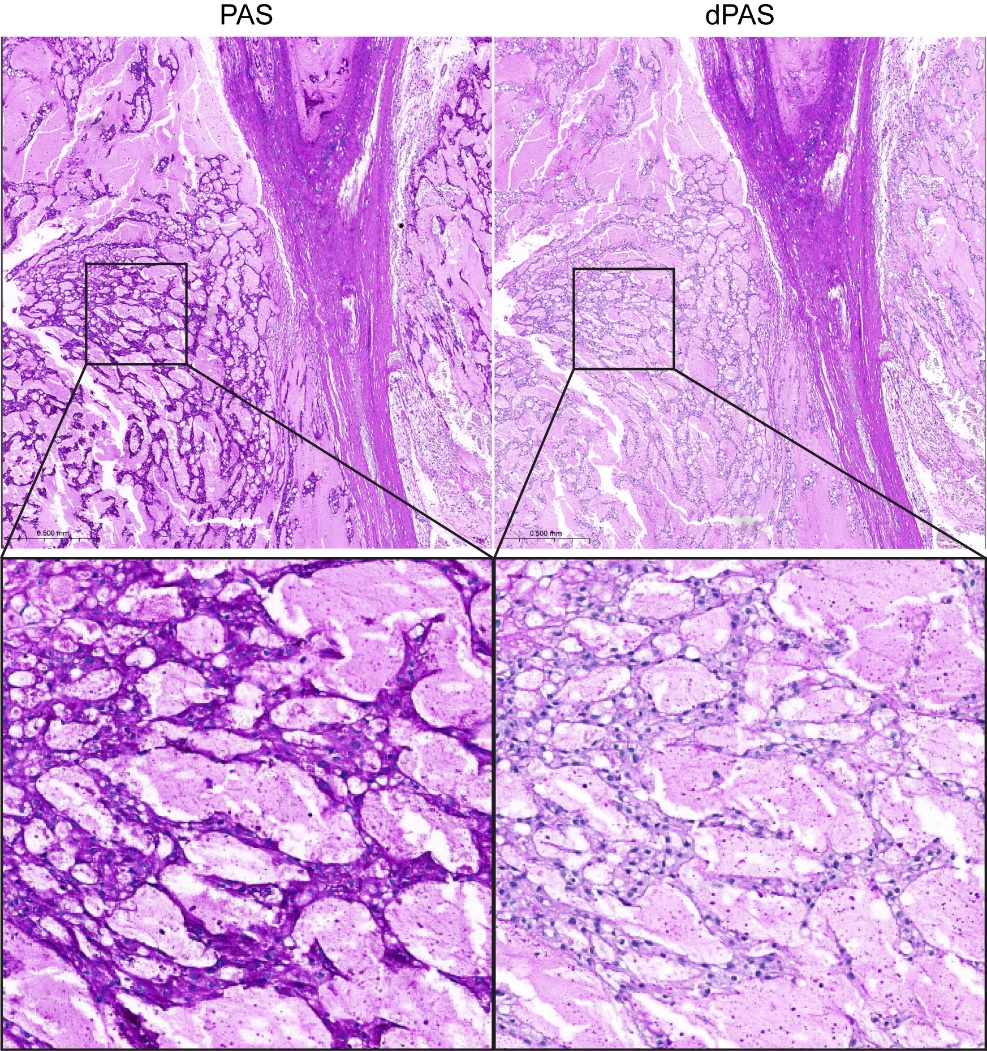

**Supplementary Figure 1. PAS/dPAS staining shows cytoplasmic glycogen.** PAS and dPAS staining from corresponding areas in the same sample on consecutive FFPE slides, showing the loss of glycogen after digestion with diastase.

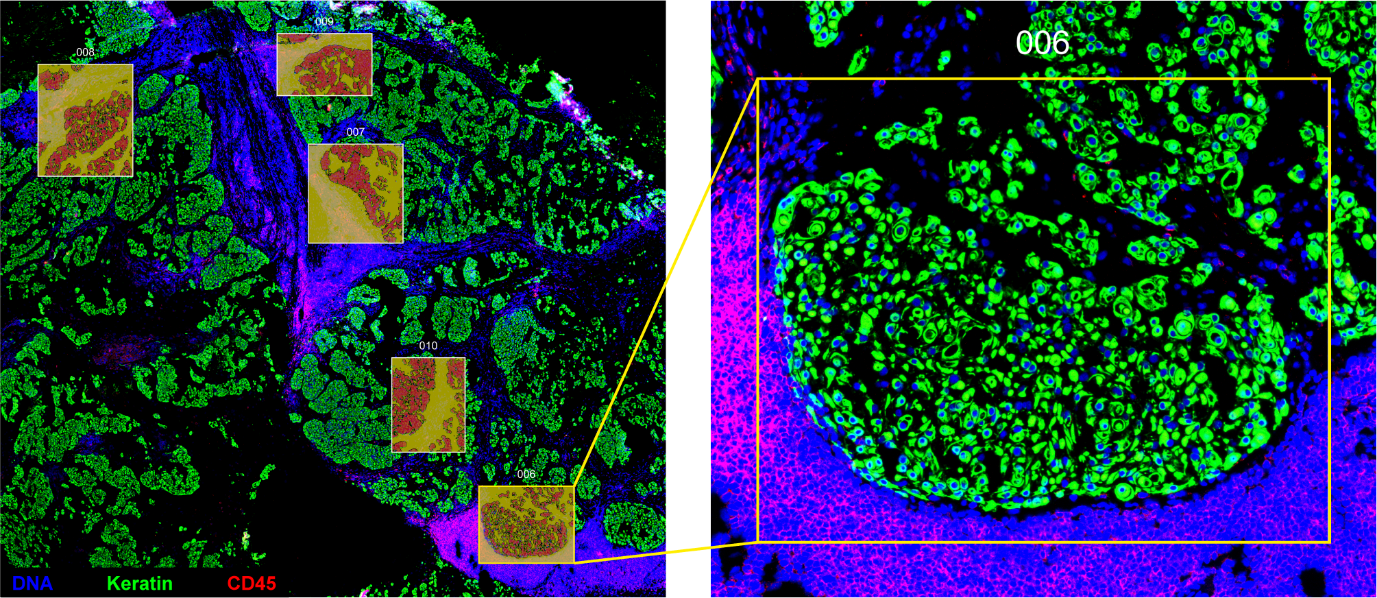

**Supplementary Figure 2. GeoMx digital spatial profiling ROI selection**. Representative immunofluorescence image of a chordoma, annotated with the selected ROIs for the GeoMx experiment. The ROIs were segmented based on keratin (green), thereby generating tumor and stroma segments. Stroma segments contained all other captured cells (DNA in blue), including immune cells (CD45 in red).

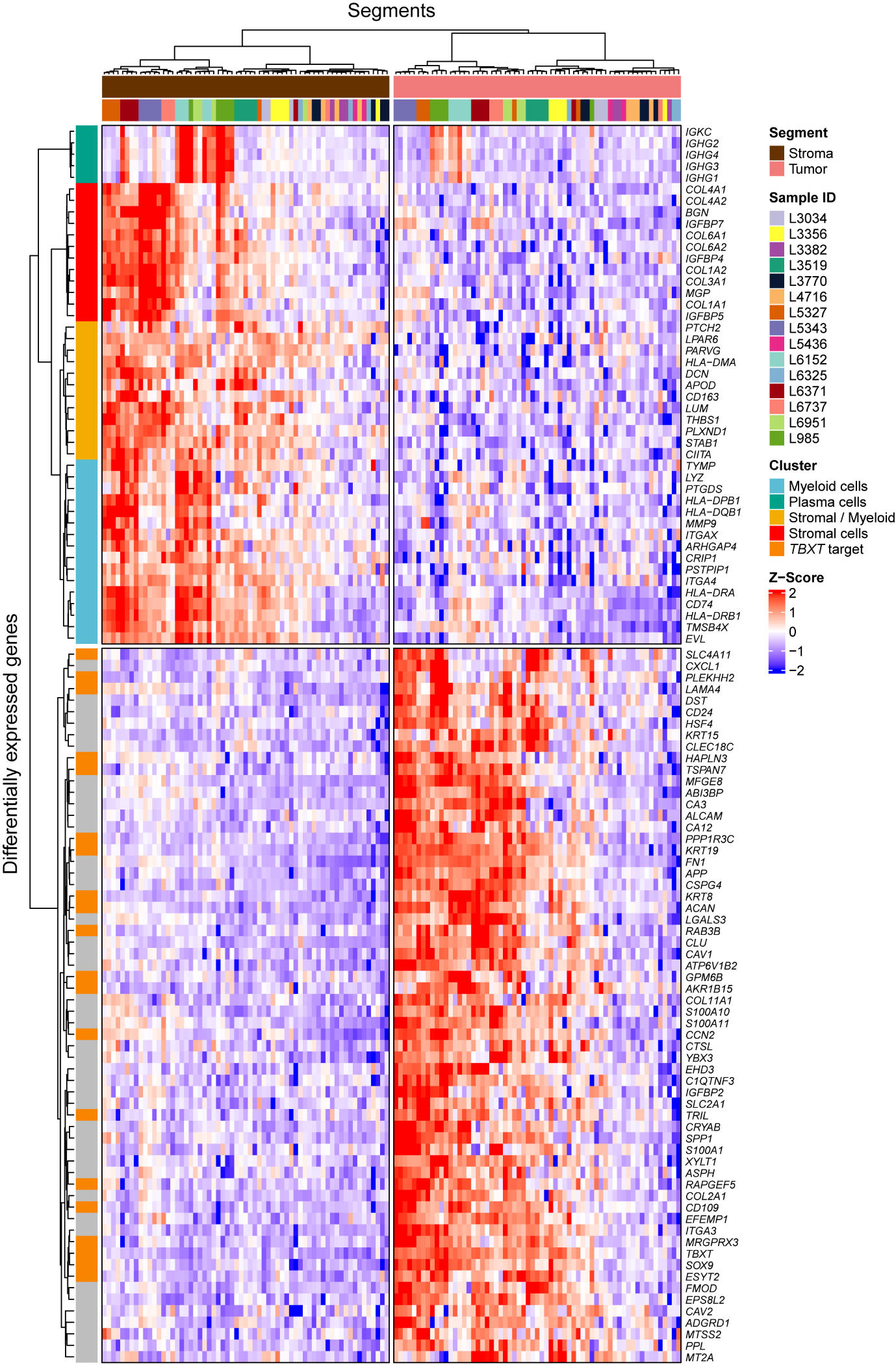

**Supplementary Figure 3. DEGs in tumor and stroma segments.** Heatmap displaying the DEGs comparing tumor with stroma segments, also presented as volcano plot in **Figure 1a**. Segments are annotated for their tissue compartment and sample ID. Genes are separated by their enriched segment. In addition, stroma DEGs are annotated for cell type clusters and tumor DEGs are annotated if they are a target of *TBXT* (1). Differential gene expression analysis was performed as described in **Figure 1**.

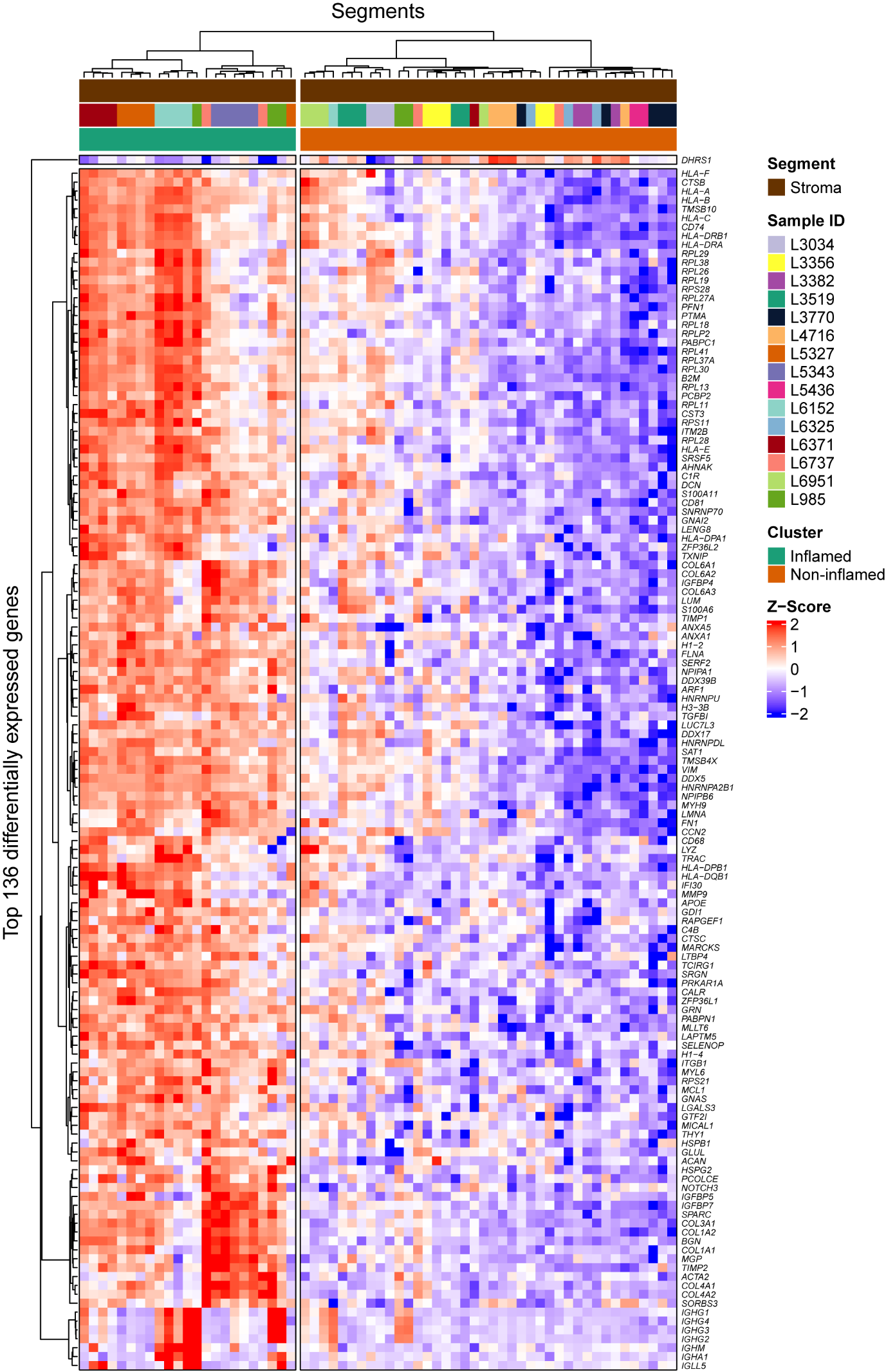

**Supplementary Figure 4. DEGs in inflamed and non-inflamed stroma segments.** Heatmap displaying the DEGs comparing inflamed with non-inflamed stroma segments, also presented as volcano plot in **Figure 1d**. Segments are annotated for their tissue compartment, sample ID and cluster. Genes are separated by their enriched cluster. Differential gene expression analysis was performed as described in **Figure 1**.

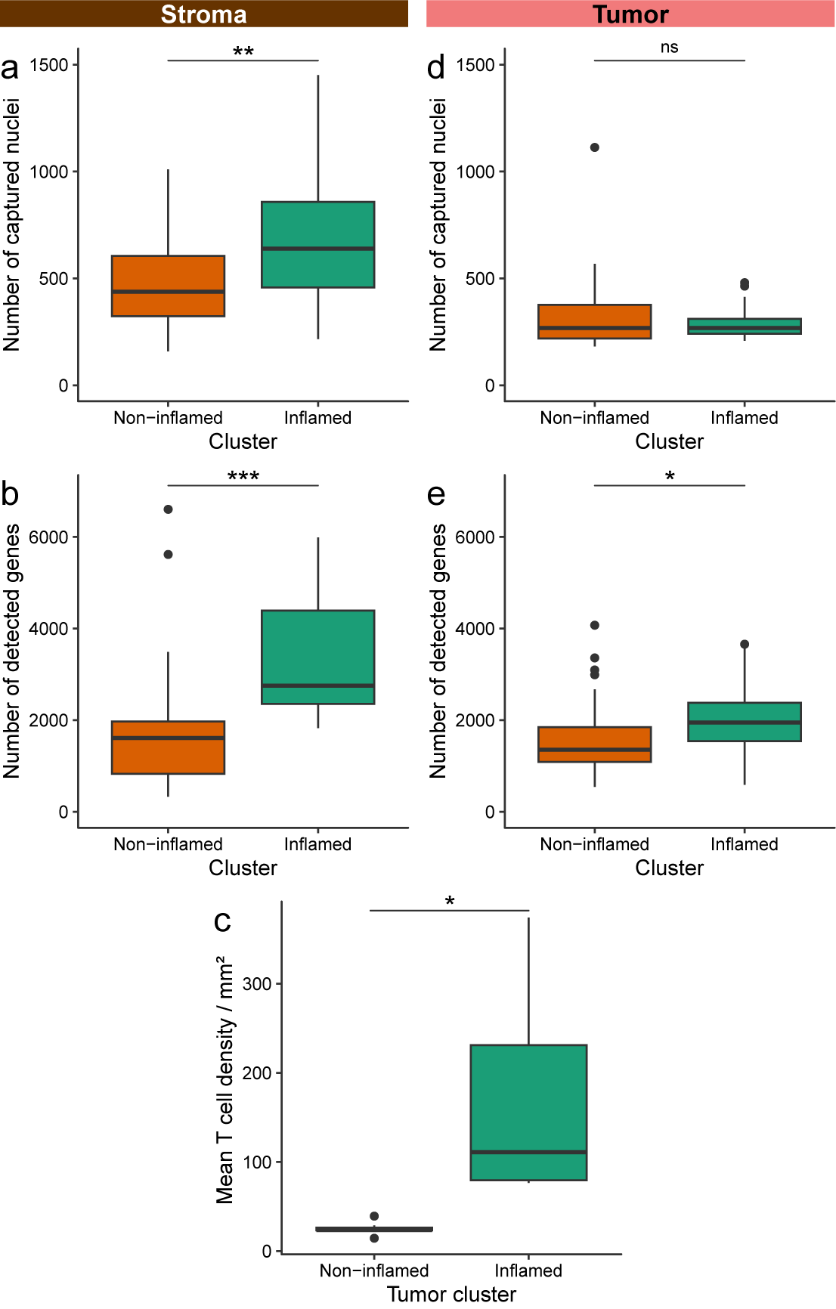

**Supplementary Figure 5. Feature comparisons between inflamed and non-inflamed segments**. a+b) Boxplots presenting the number of captured nuclei (a) and the number of detected genes (b) per stroma segment, separated by the assigned clusters. Statistical significance was assessed by unpaired Student’s t-tests. * = *P* < 0.05; ** = *P* < 0.01;*** = *P* < 0.001; ns = not significant. c) Boxplots presenting the mean T cell density / mm^2^, as previously determined by imaging mass cytometry (2), for the assigned tumor clusters. Statistical significance was assessed by performing an unpaired Student’s t-test. d+e) Boxplots presenting the number of captured nuclei (d) and the number of detected genes (e) per tumor segment, separated by the assigned clusters. Statistical significance was determined by unpaired Student’s t-tests.

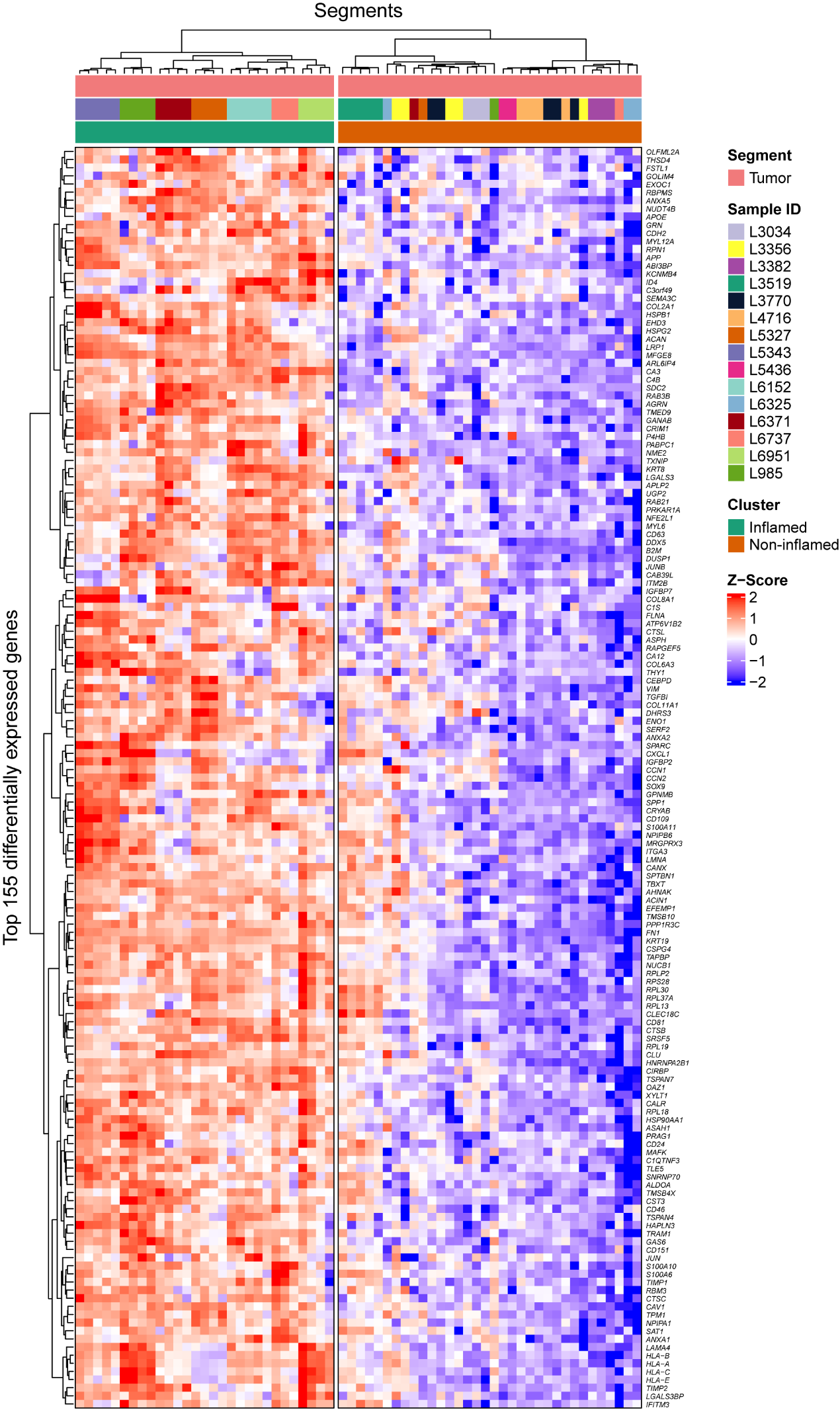

**Supplementary Figure 6. DEGs in inflamed and non-inflamed tumor segments.** Heatmap displaying the DEGs comparing inflamed with non-inflamed tumor segments, also presented as volcano plot in **Figure 1f**. Segments are annotated for their tissue compartment, sample ID and cluster. Differential gene expression analysis was performed as described in **Figure 1**.

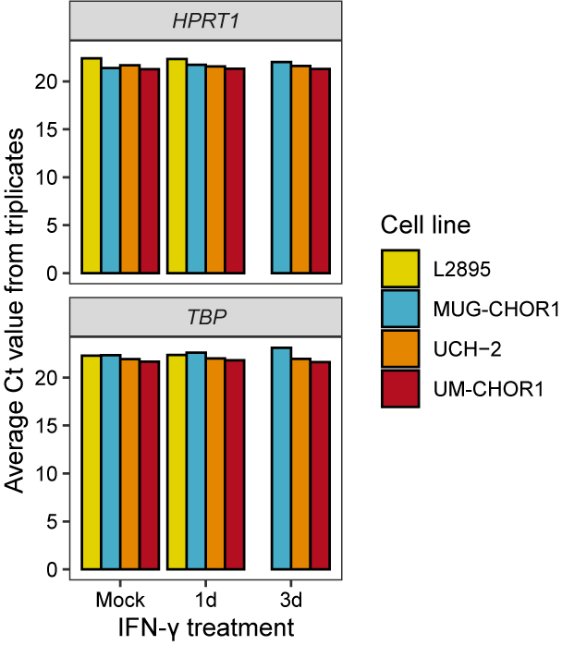

**Supplementary Figure 7. Housekeeping genes remain stable after IFN-γ treatment**. Average Ct values for *HPRT1* and *TBP* from technical triplicates for each chordoma cell line and treatment condition.

**Supplementary Table 1.** Overview of the studied samples. RT = radiotherapy.

| **Sample ID** | **Patient ID** | **Anatomical location** | **Sample type** | **Neoadjuvant RT** | **GeoMx** | **MALDI** |
| --- | --- | --- | --- | --- | --- | --- |
| L985 | P1 | Sacrum | Recurrence |  | Yes | Yes |
| L2038 | P1 |  | Recurrence |  |  | Yes |
| L4336 | P1 |  | Recurrence |  |  | Yes |
| L3034 |  | Sacrum | Primary |  | Yes | Yes |
| L3356 |  | Sacrum | Primary |  | Yes |  |
| L3382 |  | Clivus | Primary |  | Yes | Yes |
| L3519 |  | Sacrum | Primary |  | Yes | Yes |
| L3770 |  | Sacrum | Primary |  | Yes | Yes |
| L4716 |  | Sacrum | Primary |  | Yes | Yes |
| L5327 |  | Sacrum | Primary | Yes | Yes | Yes |
| L5343 | P2 | Sacrum | Primary |  | Yes | Yes |
| L6448 | P2 |  | Recurrence |  |  | Yes |
| L5436 |  | Mobile Spine (Lumbar) | Primary |  | Yes | Yes |
| L6152 |  | Left flank soft tissue | Recurrence |  | Yes | Yes |
| L6325 |  | Sacrum | Primary | Yes | Yes | Yes |
| L6371 |  | Sacrum | Primary |  | Yes | Yes |
| L6737 |  | Clivus | Primary |  | Yes | Yes |
| L6951 |  | Sacrum (Coccyx) | Primary | Yes | Yes |  |

**Supplementary Table 2.** Matrix deposition details.

| **Parameter** | **Setting** |
| --- | --- |
| Nozzle temperature (C) | 60 |
| Passes | 20 |
| Flowrate (μL / min) | 80 |
| Velocity (mm / min) | 2000 |
| Track spacing (mm) | 3 |
| Pattern | CC |
| Nitrogen pressure (psi) | 10 |
| Drying time between passes (s) | 30 |

**Supplementary Table 3**. Molecular ions detected by FTICR used for calibration of the HRAM profiling data.

| **Compound** | **Accurate mass** |
| --- | --- |
| LPE(18:1) | 478.2939 |
| LPE(22:6) | 524.2783 |
| GSSG | 611.1452 |
| Heme | 615.1700 |
| PE(36:2) | 742.5392 |
| PE(38:4) | 766.5392 |
| PS(38:4) | 810.5290 |
| PI(38:4) | 885.5498 |

**Supplementary Table 4**. Molecular ions commonly detected by MALDI-MSI when using NEDC as matrix, used here for internal mass calibration.

| **Compound** | **Accurate mass** |
| --- | --- |
| Taurine | 124.006841 |
| Phosphoethanolamine | 140.01187 |
| L-Glutamic acid | 146.045334 |
| F6P | 259.022446 |
| Inosine | 267.073494 |
| Linoleic acid | 279.232953 |
| Oleic acid | 281.248603 |
| Stearic acid | 283.264253 |
| Arachidonic acid | 303.232953 |
| Glutathione | 306.076532 |
| AMP | 346.055811 |
| ADP | 426.022144 |
| LPE(18:1) | 478.293915 |
| LPE(22:6) | 524.2783 |
| GSSG | 611.145272 |
| Heme | 615.170018 |
| PE(36:2) | 742.53923 |
| PE(38:4) | 766.53923 |
| PS(38:4) | 810.52906 |
| PI(38:4) | 885.549856 |
